# Residual barcode-like sequences and barcode-associated artifacts after Dorado and Barbell preprocessing of metagenomic reads

**DOI:** 10.64898/2026.11.05.764534

**Authors:** Timo Dinse, Vincent Robin Engländer, Silas Metz, Hubert Schrezenmeier, David Alexander Christian Messerer

## Abstract

**Background:** Residual barcode-like sequences after preprocessing are a quality-control concern in multiplexed Oxford Nanopore Technologies (ONT) sequencing data, particularly in low-biomass, host-dominated samples such as blood. Using blood datasets spiked with a microbial mock community and prepared with the Native Barcoding Kit 96 V14 (SQK-NBD114.96), we compared Dorado 1.1.1, Dorado 2.0.0, and Barbell 0.3.2.

**Results:** Dorado demultiplexing reduced barcode-positive reads to approximately 2% of reads, with internal barcode-like matches in 0.17–0.19% of reads. Barbell 0.3.2 applied to untrimmed Dorado basecalls yielded an approximately 50-fold lower detector-defined internal-match rate (0.0035–0.0037%) while retaining 91.2–91.8% of input reads. In the NCBI core_nt release of 12 September 2026, the focused screen identified complete, ungapped 24 nt identity for 64 of the 96 barcodes (66.7%), comprising 505 distinct barcode-accession pairs across 482 accessions and 195 exported TaxID groups of mixed taxonomic ranks. Records annotated as rRNA genes, chromosomes/genomes, and plasmids accounted for 35.6%, 23.2%, and 22.4% of the pairs, respectively. The barcode was located within its native ligation flanks in 224 pairs (44.4%), including 170 (33.7%) with the complete Dorado barcode search window and 3 (0.6%) with the full adapter construct.

**Conclusion:** Barbell combined 91–92% read retention with lower detected internal barcode signals. After demultiplexing, barcode-like sequences remained in sample data. Moreover, kit barcode sequences, often within their native ligation flanks, were detected in reference records. Both findings call for careful preprocessing, reference inspection, and downstream validation. Further research should determine whether barcode-associated artifacts in reads and reference records affect taxonomic classification in clinical metagenomic workflows.

## Background

Metagenomic sequencing enables culture-independent detection of microbial DNA from complex clinical samples and has been investigated for rapid pathogen detection in host-dominated clinical matrices, such as lower respiratory tract samples and plasma [1, 2]. High-throughput applications require multiplexing, in which samples are pooled after barcode ligation, sequenced together, and separated computationally, making accurate barcode recognition and post-processing removal a critical step [3, 4].

Barcode-associated artifacts arise during library preparation and bioinformatic processing [3–5]. Ligation can produce barcode-adapter constructs, barcode-barcode products, adapter dimers, and chimeras, and oligonucleotide synthesis errors can alter or truncate barcodes. Such artifacts appear as short reads or as reads in which biological sequence is interrupted by an internal barcode-like segment (Fig. 1A). During demultiplexing, sequencing errors, partial barcodes, and barcodes outside the end-anchored search window can prevent recognition, so that reads remain unclassified, are misassigned, or keep barcode remnants after trimming [5]. Dedicated tools remove such residues by matching known adapter sequences or by inferring them de novo [6, 7]. Residual barcodes can distort read-length statistics, mapping, and taxonomic classification, and can contribute to sample crosstalk [3–5], particularly in low-biomass, host-dominated samples [2, 8]. For k-mer classifiers such as Kraken 2, barcode remnants and barcode–insert junctions can add misleading evidence (Supplement S1). Whether this causes misclassification can only be tested with the classifier itself [5]. Public sequence databases also contain Oxford Nanopore Technologies (ONT) barcode and adapter sequences in GenBank records [9, 10], so reference records should be checked for the barcodes used in a given workflow.

**Figure 1.**
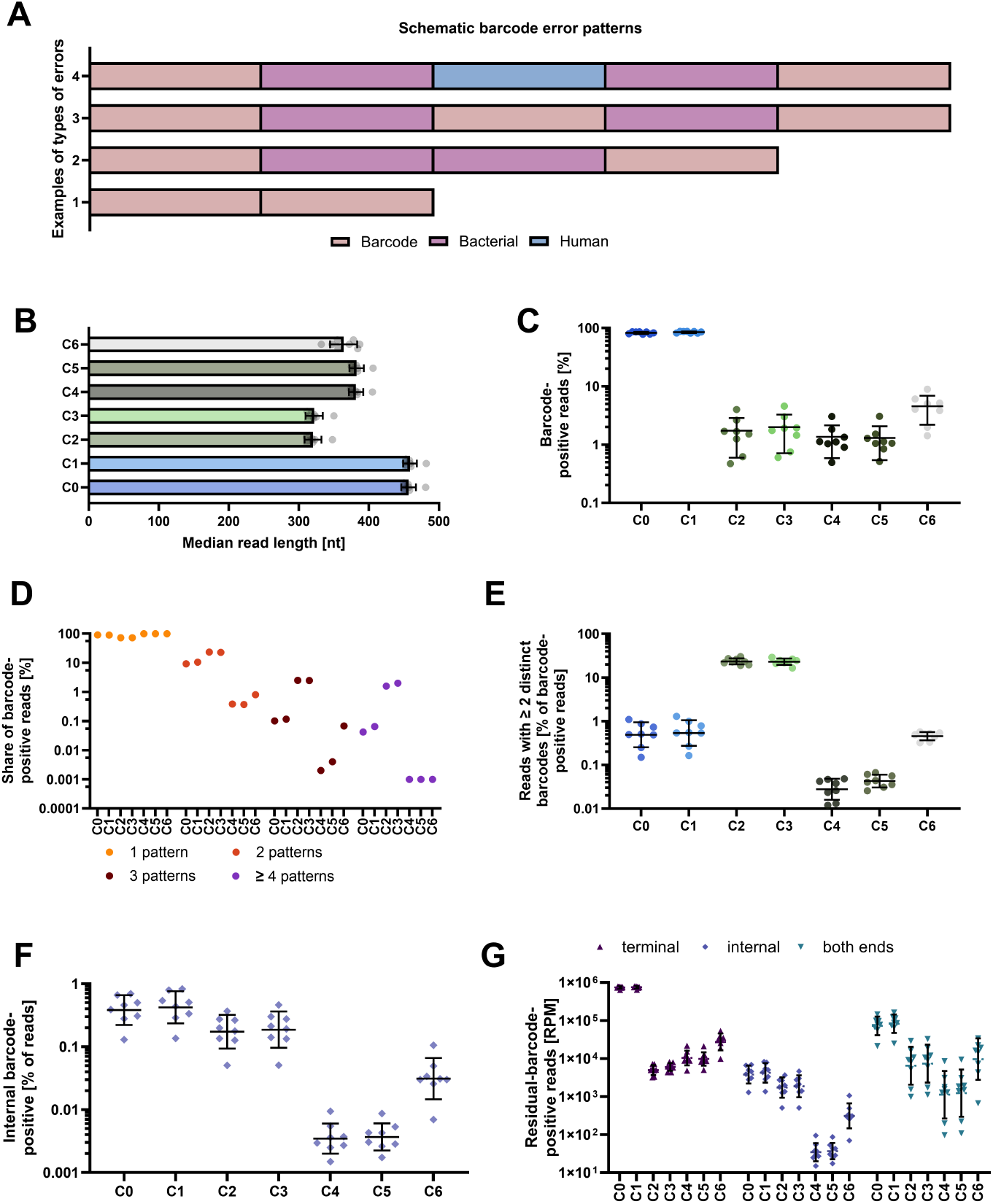
Barcode-associated artifacts and read-level measurements across preprocessing conditions C0–C6. (A) Schematic examples of a barcode-only sequence (Error 1), bacterial sequence still flanked by barcodes after trimming (Error 2), an internal barcode between bacterial segments (Error 3), and mixed bacterial/human sequence between terminal barcodes (Error 4). The examples are illustrative and are not molecular structures validated by the residual scan. (B) Median read length: bars show the mean of eight experiment-specific medians; symbols are the eight per-experiment medians, and error bars show the SD. (C) Percentage of all output reads with at least one approximate barcode match under the Methods edit budget. Symbols are the eight per-experiment values, lines the mean, and whiskers the SD. The y-axis uses a base-10 logarithmic scale; tick labels report percentages, not log10-transformed values. (D) Share of barcode-positive reads with one, two, three, or at least four distinct barcode-orientation patterns, averaged over eight experiments. A pattern is one barcode ID in one orientation; repeated occurrences of the same pattern count once (occurrence counts in Supplement S6.3). Symbols are grouped by pattern class and show C0–C6 from left to right; for each condition, the four classes sum to 100%. The y-axis is logarithmic. Zero values (at least four patterns in C4–C6) are plotted at 0.001%. (E) Percentage of barcode-positive reads matching more than one distinct barcode ID. This is a detector-defined endpoint, not directly measured inter-sample crosstalk. (F) Percentage of all output reads with a detected internal barcode-like match. (G) Mutually exclusive location classes per million output reads: internal if any detected match has `start` > 80 and `read_length` − `end` > 80; otherwise both-ends if the left and right terminal tests are satisfied; otherwise terminal. Coordinates are zero-based and end-exclusive. A single match on a short read may satisfy both end tests. Matches are all non-overlapping occurrences within the edit budget (Methods). In E–G, symbols are the eight experiments, central lines are geometric means, and whiskers span geometric mean multiplied/divided by GSD on logarithmic axes. GSD describes dispersion, not a confidence interval. Each condition uses its own output population; C6 is the selected subset returned from C3 by sequential Barbell processing. The geometric means of the location classes are not additive and do not sum to the barcode-positive rate in C. Descriptive statistics are in Supplementary Table S4; per-experiment read yields and pooled read-length distributions are in Supplementary Figure S1. GSD, geometric standard deviation; ID, identifier; SD, standard deviation.

Dorado, the ONT basecaller, combines basecalling with barcode classification, demultiplexing, and trimming, using basecalling models matched to each software release [11]. Barbell is a dedicated post-basecalling tool designed to detect and remove complex barcode-adapter patterns, including internal barcode fragments that can remain after standard demultiplexing and trimming [5]. How these tools compare in the barcode residues they leave in host-dominated metagenomic data, and whether such residues affect taxonomic classification, is not yet clear.

Here, we compare Dorado 1.1.1, Dorado 2.0.0, and Barbell 0.3.2 on eight blood datasets spiked with a microbial mock community and prepared with the Native Barcoding Kit 96 V14 (SQK-NBD114.96). We assess read retention, read-length distributions, and residual barcode-like sequences, detected with a predefined approximate-matching rule. A complementary screen tests whether the kit’s barcodes, alone or within their native constructs, also occur with complete-query identity in NCBI core_nt reference records.

## Methods

### Sample preparation and sequencing

Blood from eight healthy male volunteers (25–67 years) was obtained from the initial predonation aliquot collected during routine donation venipuncture. All donors had provided written informed consent for the research use of residual sample material. Samples were spiked with the ZymoBIOMICS Gut Micro-biome Standard (a microbial mock community consisting of 18 bacterial, 2 fungal, and 1 archaeal strains; Zymo Research) at one of five levels or left unspiked. Each of the eight experiments comprises the six barcoded samples of one donor (unspiked and five spike levels) and was treated as a replicate. DNA was extracted with an adapted Ultra-Deep Microbiome Prep protocol (Molzym) without host-DNA depletion, and libraries were prepared with the SQK-NBD114.96 kit (ONT; Supplement S2). Sequencing used a PromethION P24 device (R10.4.1 flow cells) with adaptive sampling against the human genome (hg38). MinKNOW performed live basecalling using the high-accuracy (HAC) model and classified reads with a mean Q-score ≥ 9 as pass reads. Of these, 84.3% were rejected and thus truncated by adaptive sampling (Supplement S3). All reads, including the 3.9% classified as fail, were subsequently rebasecalled without Q-score filtering.

### Basecalling and preprocessing conditions

Each experiment was processed in seven conditions (Supplement S4 and S5; Supplementary Tables S1 and S2). C0 and C1 are untrimmed basecalls from Dorado 1.1.1 (HAC model v5.0.0) and Dorado 2.0.0 (HAC v6.0.0) [11]; C2 and C3 add Dorado’s demultiplexing and trimming [11]. C4 and C5 apply Barbell 0.3.2 with its default filter to the untrimmed C0 and C1 reads [5], and C6 applies it to the complete C3 output [5, 11]. The filter keeps reads with a recognized barcode at one end or concordant barcodes at both ends, and Barbell has no unclassified output (Supplement S5.2). Comparisons are therefore between output populations: C2 and C3 include Dorado’s unclassified reads, C4–C6 only accepted reads. Dorado 2.0.0 BAM files were converted to FASTQ with SAMtools [12] (Supplement S5.3).

### Residual-barcode detection and statistics

All output reads were scanned for the 192 barcode orientations (96 barcodes and their reverse complements) with the Python `regex` package, allowing a Levenshtein distance [13] of at most 2 per 24 nucleotide (nt) barcode and counting every non-overlapping occurrence (Supplement S6.1–S6.3). Matches within 80 nt of a read end were terminal, all others internal. Reads with any internal match were classified as internal. Among reads without internal matches, those with terminal matches at both ends were classified as both-ends, whereas reads with a terminal match at only one end were classified as terminal. The primary endpoint was the rate of reads with an internal match per million output reads (RPM). Planted variants (Supplement S6.6) and shuffled decoy barcodes in three experiments (Supplement S6.7; Supplementary Table S3) tested detection and chance matching. Read Q-scores were summarized with NanoStat [14] (Supplement S6.4).

Rates spanning orders of magnitude were summarized across the eight experiments as geometric mean ×/÷ geometric standard deviation (GSD), a unitless fold factor, and other values as mean ± standard deviation (SD; Supplement S7.3; Supplementary Table S4). Read-level outcomes were described without testing (Supplementary Table S5). Internal rates were compared with the paired two-sided Wilcoxon signed-rank test without multiplicity adjustment. Fold-changes are geometric means of the per-experiment ratios (Supplementary Table S6; Supplement S8).

### Focused ONT barcode-reference sequence analysis

To test whether the kit’s barcodes also occur in reference records, the 96 barcodes were searched alone (B24) and within three native constructs defined in Dorado 1.1.1, which add the ligation flanks (K40), the complete Dorado barcode search window (K46), or the sequencing adapter (K68) (Supplement S9.1). BLASTn [15] (`blastn-short`) searched the NCBI core_nt release of 12 September 2026, and only ungapped alignments with 100% identity over the complete query were retained (Supplement S9.2–S9.3; Supplementary Table S7). Matches were counted as distinct barcode-accession pairs and grouped by NCBI TaxID and record title (Supplement S9.4–S9.5).

## Results

### Read yield and length characteristics

Dorado demultiplexing (C2/C3) retained 99.94% of the untrimmed reads, of which 6.0% (C2) and 5.9% (C3) were in its unclassified output. Barbell retained 91.8% (C4) and 91.2% (C5) of the input reads. Median read length was 457–459 nt in untrimmed output, 320–322 nt after Dorado, and 382–383 nt after Barbell (Fig. 1B). Dorado output had more short reads and a broader length distribution than Barbell output (Supplementary Figure S1; Supplement S10.1). Applied after Dorado (C6), Barbell returned only 0.74% of the C3 reads, consistent with its filter: after Dorado trimming, only 2.0 ± 1.3% of C3 reads still contained any barcode-like match (Fig. 1C).

### Residual barcode burden

Barcode-positive reads, with at least one match anywhere in the read, made up 82.9 ± 3.4% (C0) and 85.1 ± 2.8% (C1) of untrimmed output. After processing, they made up 1.7 ± 1.1% (C2), 2.0 ± 1.3% (C3), 1.4 ± 0.8% (C4), and 1.3 ± 0.8% (C5) (Fig. 1C). Dorado 2.0.0/HAC v6.0.0 did not lower this rate, which was slightly higher than with Dorado 1.1.1/HAC v5.0.0 in both the untrimmed and demultiplexed pairs. In the C6 subset, 4.5 ± 2.4% of the remaining reads were positive.

The positive reads differed in composition. After Dorado demultiplexing, 27% of them carried two or more distinct barcode-orientation patterns, compared with 0.4% after Barbell (Fig. 1D; Supplement S6.3). The share of barcode-positive reads matching more than one barcode ID rose from 0.49% (C0) and 0.54% (C1) to 23.4% (C2) and 23.1% (C3), but was only 0.028% (C4) and 0.043% (C5) after Barbell (Fig. 1E). This apparent increase after Dorado demultiplexing reflects a change in the denominator rather than an increase in multi-ID reads. Per million output reads, multi-ID reads changed little (4,050 RPM in C0 versus 3,322 RPM in C2), whereas barcode-positive reads fell from approximately 829,000 to 14,000 RPM, most likely because trimming removed mainly single terminal barcodes. Barbell does not output reads with discordant barcodes, reducing multi-ID reads to 3.3 RPM (C4) and 4.9 RPM (C5).

### Terminal versus internal residual barcodes

Internal matches differed most between configurations (Fig. 1F, G). They occurred in 0.38% (C0) and 0.42% (C1) of untrimmed reads (geometric means; Supplementary Table S4), in 0.17% (C2) and 0.19% (C3) after Dorado demultiplexing, and in 0.0035% (C4) and 0.0037% (C5), or 35–37 RPM, after Barbell. This approximately 50-fold reduction occurred in all eight experiments and for both Dorado versions (C2 versus C4: geometric-mean fold-change 50.0×, range 33–74; C3 versus C5: 50.7×, range 29–70; Wilcoxon p = 0.0078 for both, the minimum attainable with eight pairs; Supplementary Table S6). Terminal-only matches dominated untrimmed output (742,441 RPM in C0 and 751,724 RPM in C1, about three of four reads) and fell to 4,945 RPM in C2 and 5,996 RPM in C3 (GSD 1.36 and 1.26, i.e., experiments typically within 1.4-fold of the geometric mean), but were higher after Barbell (10,368 RPM in C4 and 9,766 RPM in C5; 2.10- and 1.63-fold).

The experiments differed considerably, least for terminal matches in untrimmed output (GSD 1.07) and most for both-ends matches (GSD 1.76 in C0, up to 4.22 in C4). The paired analysis accounts for this variability (Supplement S10.2). Shuffled decoy barcodes produced internal matches at 0–2.3 RPM per decoy set in C0–C5 and none in C6 (Supplementary Table S3). In the same experiments, this chance background was at most 0.3% of the internal rate after Dorado demultiplexing and at most 7.8% (C4) and 5.1% (C5) of the rate after Barbell. All rates are reported without subtracting it.

### Matching ONT barcode sequences in reference records

Under the strict complete-query criterion, 64 of the 96 barcodes (66.7%) matched reference records, in 505 distinct barcode-accession pairs across 482 accessions and 195 exported TaxID groups (Fig. 2A; Supplementary Table S7). NB08 matched the most accessions (84), followed by NB03 (37) and NB04 (35; Fig. 2B). Nineteen barcodes matched a single accession (Supplementary Table S8). The TaxID groups were of mixed rank (87 bacterial, 62 eukaryotic, and 46 viral). Those with the most pairs were *Plasmodium berghei* ANKA (69), uncultured *Muribaculaceae* bacterium (31), *Escherichia coli* (22), and *Klebsiella pneumoniae* (19). Records annotated as rRNA genes, chromosomes/genomes, and plasmids contributed 35.6%, 23.2%, and 22.4% of the pairs (Fig. 2C; Supplement S9.4–S9.5).

**Figure 2.**
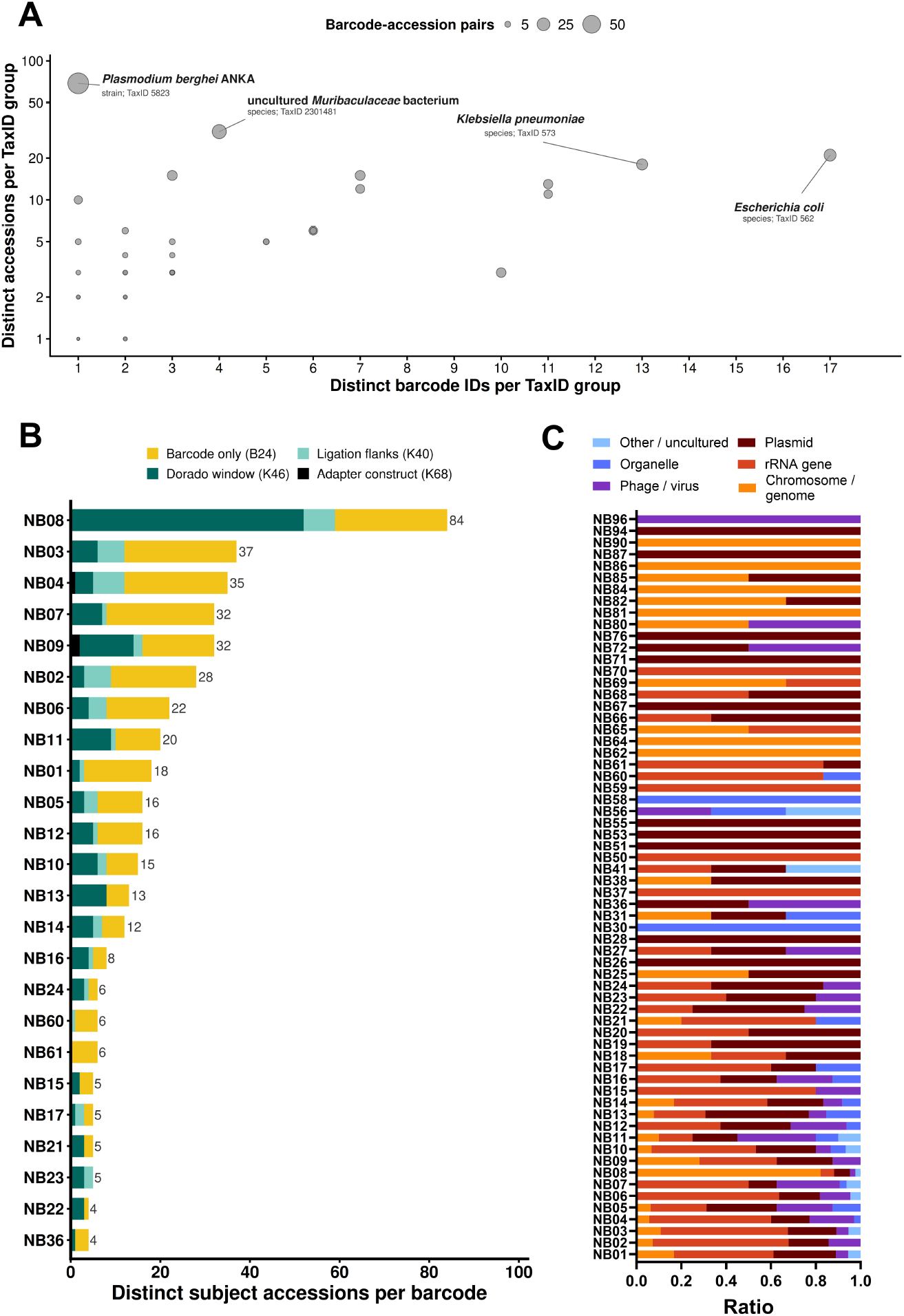
Focused ONT SQK-NBD114.96 case study: complete barcode matches in reference records and their native construct context. Searches used the NCBI core_nt release of 12 September 2026. A match requires complete, ungapped identity over the whole query: `pident` = 100%, `mismatch` = 0, `gapopen` = 0, query endpoints 1 and `qlen`, `length` = `qlen`, and `abs(send - sstart)` + 1 = `qlen`. (A) Barcode-only (B24) matches grouped by the 195 exported TaxIDs: x, number of distinct barcode IDs; y, number of distinct subject accessions (logarithmic scale); bubble area, number of barcode-accession pairs. Coincident points overlap. Labels identify the four groups with the most pairs, with scientific names, ranks, and TaxIDs from NCBI Taxonomy (9 October 2026). All exported groups are kept without species-level collapse and include bacterial, eukaryotic, and viral entries of mixed rank (Supplement S9.5). (B) Distinct subject accessions per barcode for the 24 barcodes with more than three accessions (434 of 505 pairs), sorted by count with barcode ID as the tie-breaker. Bar length is the number of barcode-only accessions; colored segments divide these pairs by the longest native construct matched at the same record locus, as defined in Dorado 1.1.1: barcode only (B24), ligation flanks (K40; AAGGTTAA + barcode + CAGCACCT), Dorado barcode search window (K46; ATTGCTAAGGTTAA + barcode + CAGCACCT), or adapter construct (K68; LSK110 adapter joined to the K46 window). Barcodes with three accessions (NB18, NB19, NB27, NB31, NB38, NB41, NB56, NB66, NB69, and NB82; 30 pairs) and with two accessions (NB20, NB25, NB28, NB64, NB65, NB68, NB72, NB80, NB81, NB85, and NB86; 22 pairs) are not shown. The 19 barcodes with a single accession are listed in Supplementary Table S8; 32 barcodes had no match. (C) Record-title category composition of the barcode-only pairs of each of the 64 hit-bearing barcodes, ordered by barcode ID, as horizontal stacked bars on a ratio scale from 0 to 1 (n = 505 pairs). Each pair contributes once to one title-keyword category; the chromosome/ genome category is not restricted to bacteria. HSP-weighted composition (n = 1,390 HSPs) and the composition of the construct panels are given in Supplement S9.4. Counts are descriptive and are not false-positive probabilities. See Supplement S9 and Supplementary Tables S7 and S8. These matches motivate reference inspection; they do not classify records as contaminated or link their origins to the reads from the sequenced blood samples. HSP, high-scoring segment pair; LSK110, Ligation Sequencing Kit 110; NCBI, National Center for Biotechnology Information; ONT, Oxford Nanopore Technologies; TaxID, NCBI Taxonomy identifier.

Many matches extended into kit sequence (Fig. 2B). In 224 pairs (44.4%; 42 barcodes), the record carried the barcode with both native ligation flanks (K40), including 170 pairs (33.7%) with the complete Dorado barcode search window (K46) and 3 (0.6%) with the full adapter construct (K68). Flanked matches were common in organelle, phage/virus, chromosome/genome, and plasmid records but less common in rRNA gene records (44 of 180 pairs), and record sequences retrieved from NCBI confirmed every construct call (Supplement S9.6). Because a K68 match required identity over all 68 nt, and the first adapter bases are often missing or miscalled in native-barcoded reads, library-derived sequence in a record would mostly match only K40 or K46, so the three K68 matches probably underestimate adapter-derived sequence (Supplement S9.1).

## Discussion

This study makes two observations for the SQK-NBD114.96 kit. First, barcode-like sequences remain in reads after processing, and Dorado and Barbell differ mainly in where these residual sequences lie. Second, the kit’s barcodes also occur in reference records, often within native kit flanks. Terminal remnants and internal barcode or adapter patterns have been described in multiplexed ONT data, and internal patterns can indicate chimeric ligation products [3–5]. Such sequences may add spurious k-mers and short local alignments for downstream classifiers [5].

After Dorado demultiplexing and after Barbell, a similarly small fraction of reads remained barcode-positive, but the kind of residue differed. Under Dorado, 38 ± 16% (C2) and 39 ± 18% (C3) of the positive reads had matches near one read end alone, and nearly a quarter matched two or more barcode IDs. Under Barbell, almost nine in ten (87 ± 9% in C4 and 85 ± 10% in C5) had matches near one read end alone. Dorado searches for barcodes at the read ends, trims them, and writes almost every read, including reads with internal or discordant barcodes. Barbell discards reads whose barcode configuration it does not accept, which most likely explains both its approximately 50-fold lower internal rate and its loss of 8–9% of the reads. Its smaller share of short reads may reflect trimming, selective retention, or both. The residues that Barbell leaves are mainly remnants at read ends, and its terminal-only rate was higher than Dorado’s, so neither configuration was better on every endpoint. Terminal remnants add kit sequence to a read end, whereas internal barcodes can join unrelated inserts. Running Barbell after Dorado processing (C6) returned only a small subset enriched for residual barcodes, because Dorado had already removed most of the barcodes that Barbell needs to accept a read.

The decoy scan shows that chance matching explains at most a small part of the internal matches after Barbell, and subtracting it would widen rather than narrow the difference. Dorado 2.0.0 with HAC v6.0.0 did not reduce the residual burden, and detection was even slightly higher than with Dorado 1.1.1. Better basecall accuracy could expose previously missed barcodes, whereas better barcode recognition could reduce their survival after trimming, and the data cannot separate these effects. The median predicted read Q-scores were lower under HAC v6.0.0 in all eight experiments (Supplement S6.4), but they are model-specific calibrations, not measured accuracy. Because basecalling models keep improving [16, 17] and release and model changed together here, version comparisons based on sequence similarity carry this confound.

The reference screen assessed the occurrence of workflow-relevant barcodes in reference records, with a focus on sequence matches. Public databases contain barcode and adapter contamination, including Nanopore-derived footprints [5, 9], and 44% of the matching pairs carried the barcode within its native ligation flanks, consistent with such footprints. Matches without native flanks, which dominated rRNA gene records, can also derive from kit sequence, because ONT’s PCR, rapid, and 16S barcoding kits use the same 24 nt barcodes with other flanks [11], and a single differing flank base or a record ending at the barcode also prevents a construct match. Kraken 2 classifies reads by 35 nt k-mers by default, which a bare barcode cannot fill but the 40 nt and longer constructs can, so such records in a classifier database could give reads with the same kit sequence misleading hits (Supplement S1). Whether this affects taxonomic classification, 16S/18S profiling, or resistance-gene calling was not tested. Screening tools address contamination at database submission [18], and removing adapter contamination improves microbial genome databases [19].

This study has several limitations. The experimental design was restricted to spiked donor blood, a single barcoding kit, two Dorado release–model pairs, and Barbell’s default filter, which may limit generalizability to clinical samples. The fixed edit budget may miss divergent or truncated barcodes, whereas detected matches represent operational sequence matches rather than validated physical barcode copies, chimeras, or crosstalk events. The study included neither barcode-free libraries, which would show the detector’s background in reads without kit sequence, nor extraction or library blanks. The unspiked samples control for the spike-in but not for reagent contamination, which requires reagent-only negative controls in low-biomass studies [8, 20]. Barcode-assignment accuracy was not measured, so a lower residual rate does not prove safer demultiplexing [5]. The between-experiment offset could not be attributed to donor, DNA input, or sequencing run (Supplement S10.2), and the paired test has limited resolution with eight experiments. Finally, we did not test whether fewer internal matches lead to fewer false-positive species calls, for example with Kraken 2 [21].

## Conclusion

In eight spiked blood datasets prepared with the SQK-NBD114.96 kit, Barbell 0.3.2 applied to untrimmed Dorado output retained 91–92% of reads and had an approximately 50-fold lower internal barcode-like match rate than Dorado demultiplexing, at the cost of more terminal remnants. Dorado 2.0.0 did not reduce the residual burden compared with Dorado 1.1.1. The kit’s barcodes also occur in reference records, often within native kit flanks. Residual barcodes therefore deserve attention both in reads and in reference databases. Classifier-level validation with independently verified sample assignments is needed before a processing recommendation for clinical metagenomics can be made [5].

## Supporting information

supplementary_source_data.zip

## List of abbreviations

**Abbreviation Full term**

AI: artificial intelligence
BAM: Binary Alignment/Map format
BLASTn: nucleotide Basic Local Alignment Search Tool
DNA: deoxyribonucleic acid
FASTQ: text-based sequencing-read format containing nucleotide sequences and quality scores
GSD: geometric standard deviation (multiplicative dispersion factor)
HAC: high-accuracy basecalling model
hg38: human reference genome assembly GRCh38
HSP: high-scoring segment pair (BLAST alignment)
ID: identifier
LSK110: Ligation Sequencing Kit 110 (Oxford Nanopore Technologies)
NCBI: National Center for Biotechnology Information
nt: nucleotides
ONT: Oxford Nanopore Technologies
Q-score: Phred quality score
RPM: reads per million
rRNA: ribosomal ribonucleic acid
SD: standard deviation
SQK-NBD114.96: Native Barcoding Kit 96 V14 (Oxford Nanopore Technologies)
TaxID: NCBI Taxonomy identifier

## Ethics approval and consent to participate

All experiments were conducted in accordance with the ethical standards of the Declaration of Helsinki. The Local Independent Ethics Committee of Ulm University waived the requirement for formal ethical approval (10 July 2023), as all blood samples were obtained from predonation sampling material collected from healthy human volunteers who donated blood. All participants had provided written informed consent for the use of residual material for research purposes, and no additional venipuncture was performed for this study.

## Consent for publication

Not applicable.

## Authors’ contributions

Conceptualization: TD and DM; Data curation: TD and DM; Formal analysis: TD; Funding acquisition: DM; Investigation: TD and VE; Methodology: TD and SM; Project administration: TD, HS, and DM; Software: TD and SM; Supervision: DM; Validation: TD and DM; Visualization: TD and DM; Writing – Original Draft: TD and DM; Writing – Review & Editing: TD, VE, SM, HS, and DM.

## Availability of data and material

Additional data are available in the Supplement. The scripts used for the analyses and figures and the perexperiment results underlying the figures and tables, with anonymized experiment names, are provided in Supplementary Data 1 (supplementary_source_data.zip); its contents are listed in Supplement S7.2. The scripts are also available at https://github.com/blutspende/barcode_tgs (release v1.0.1, commit fb99c9d15459) and archived at Zenodo (https://doi.org/10.5281/zenodo.23981076). Donor-level data, identifiers, raw signals, read-level sequencing files, input sequences, and acquisition and quality-control reports are excluded under the institutional data-privacy policy.

## Acknowledgements

The authors thank the blood donors who consented to the research use of residual sample material, and the staff of the German Red Cross Blood Donation Service Baden-Württemberg – Hessen and the sequencing facility for technical support. The authors also thank Daniel Kefalas and the team of the blood donation unit at IKT Ulm for their support in obtaining blood samples. The authors further thank Sarah Radecke for her contributions to the methodology and for reviewing and editing the manuscript.

## Declaration of generative AI and AI-assisted technologies in the manuscript preparation process

During the preparation of this work the authors used Microsoft Word (version 2512, Microsoft), Chat-GPT 5.6 (OpenAI), and Claude Opus 4.8 (Anthropic) for language writing and editing, improvement of readability, and coding assistance. After using these tools/services, the authors reviewed and edited the content as needed and take full responsibility for the content of the published article.

