## Supplement for "Residual barcode-like sequences and barcode-associated artifacts after Dorado and Barbell preprocessing of metagenomic reads"

#### Materials and Methods

##### S1. Kraken 2 classification parameters

Kraken 2 classifies a sequence by looking up the minimizer of each of its k-mers (with spaced-seed masking) in its database and assigns a taxon from the combined hits. An optional confidence threshold sets the fraction of a sequence's unambiguous k-mers that must map within the assigned clade. The default nucleotide database uses $k = 35$ and a minimizer length of $l = 31$, both set at database construction. A sequence of length $L$ contains $\max(0, L - k + 1)$ k-mers, and k-mers with ambiguous bases are not queried. With the default $k$, a 24 nucleotide (nt) barcode therefore contains no complete k-mer, whereas the 40, 46, and 68 nt constructs of S9.1 contain 6, 12, and 34 k-mers. Manual: `https://github.com/DerrickWood/kraken2/wiki/Manual` (accessed 11 September 2026).

##### S2. Samples, library preparation, and barcode set

*Samples and spike-in.* Blood was obtained from eight healthy male volunteers aged 25–67 years during routine blood donation visits. Study samples came from the initial predonation aliquot of the routine donation venipuncture, so no additional venipuncture was performed for the study. All donors had given written informed consent for the research use of residual sample material. Within 30 min of collection, ethylenediaminetetraacetic acid (EDTA)-anticoagulated blood was spiked with the ZymoBIOMICS Gut Microbiome Standard (Zymo Research, Irvine, CA, USA; stock concentration $3.94 \times 10^9$ cells/mL) at one of five spike levels, designated T1–T5 to keep them distinct from the processing conditions C0–C6. For the highest level (T1), 20 µL of undiluted standard was added to whole blood for a total volume of 400 µL. The four lower levels (T2–T5) were prepared by serially diluting the standard 1:10 in DNA/RNA Shield (Zymo Research) and adding each dilution in the same 20 µL volume. From each 400 µL mixture, 200 µL was used for DNA extraction, giving nominal inputs per extraction of $3.94 \times 10^7$ (T1), $3.94 \times 10^6$ (T2), $3.94 \times 10^5$ (T3), $3.94 \times 10^4$ (T4), and $3.94 \times 10^3$ (T5) microbial cells, distributed across the species according to the manufacturer's stated composition. Matched unspiked samples (T0) were processed as negative controls. Each of the eight experiments analyzed here comprises the six samples (T0–T5) of one donor, which were barcoded individually and sequenced together. Because the study addresses barcode demultiplexing and residual barcode-like sequences rather than spike-in dose-response, the experiments were treated as replicates, and spike level was not used as a covariate.

*DNA extraction and library preparation.* Genomic DNA was extracted with the Ultra-Deep Microbiome Prep kit (Molzym, Bremen, Germany) using an adapted protocol without the kit's host-DNA depletion step. Libraries were prepared according to the manufacturer's protocols for the Ligation Sequencing Kit V14 (SQK-LSK114) and the Native Barcoding Kit 96 V14 (SQK-NBD114.96; Oxford Nanopore Technologies [ONT]; Kit 14 chemistry), with DNA input volumes of 13.2–25 µL.

The kit provides 96 native barcodes (NB01–NB96) and is identified by Dorado and Barbell as `SQK-NBD114-96`. The barcode file used for residual scanning (`barcodes.csv`; ONT export format with the columns `Component`, `Forward sequence`, and `Reverse sequence`) lists each barcode in forward and reverse-complement orientation, giving 192 sequences that were searched in every read (S6).

#### S3. Basecalling models and sequencing chemistry

All sequencing runs used a PromethION P24 with R10.4.1 flow cells, Kit 14 chemistry, and a translocation speed of 400 bases per second. Adaptive sampling requires real-time basecalling, so MinKNOW basecalled the reads live with the high-accuracy (HAC) model `dna_r10.4.1_e8.2_400bps_hac@v4.3.0` and classified reads with a mean Phred quality score (Q-score) of at least 9 as pass reads (MinKNOW run reports). The offline rebasecalling kept the HAC model class:

- Dorado 1.1.1 (conditions C0 and C2; its untrimmed C0 output is also the input to C4): `dna_r10.4.1_e8.2_400bps_hac@v5.0.0`.
- Dorado 2.0.0 (conditions C1, C3, and C6; its untrimmed C1 output is also the input to C5): `dna_r10.4.1_e8.2_400bps_hac@v6.0.0`.

Recent Dorado versions require POD5 signal files, so runs recorded as FAST5 were converted first. Per experiment, all FAST5 files (pass and fail, in Exp 1 also skip) were combined and converted with the ONT pod5 tool (v0.3.39; pod5 convert fast5 <input_dir> --output <pod5_dir> --recursive). Dorado then basecalled the POD5 files of all eight experiments (option `--recursive`). The rebasecalled input of all eight experiments contained both the MinKNOW pass reads and the fail reads (2.4–11.3% fail reads per experiment, 3.9% pooled).

MinKNOW ran adaptive sampling in depletion mode against the human reference genome (hg38) with the default rejection settings. Adaptive sampling ejects a rejected molecule from the pore, which truncates its read. The MinKNOW option to skip rejected reads (`skip_rejected_reads`) was off in all runs, so rejected reads were basecalled like all other reads (MinKNOW run reports). Rejected reads were identified in the MinKNOW sequencing summaries as pass reads (`passes_filtering TRUE`) with the end reason `data_service_unblock_mux_change` (`summarize_adaptive_sampling.py`; `adaptive_sampling_pass_reads.csv` in Supplementary Data 1). They made up 70.0–88.2% of the pass reads per experiment (84.3% pooled) and had a median length of 452–480 nt, compared with 359–475 nt for the other pass reads. Rejected reads therefore formed most of the common input to all seven conditions and shaped its read-length distribution. The same table gives the read Q-scores of the pass reads (median 16.6–18.3 per experiment; S6.4).

#### S4. Software versions and invocation

All software was run under Ubuntu Linux. Versions and invocations are summarized in Table S1.

Custom analysis scripts are released in Supplementary Data 1 (`code/scripts`). They are also available at `https://github.com/blutspende/barcode_tgs` (release v1.0.1, commit `fb99c9d15459`) and archived at Zenodo (`https://doi.org/10.5281/zenodo.23981076`).

#### S5. Seven-condition preprocessing design

Eight ONT sequencing experiments were each reprocessed under the seven preprocessing conditions of Table S2 to compare residual barcode-like sequences, read yield, and read length. The experiments are referred to as Exp 1–8 here and in Supplementary Data 1. A ninth experiment was excluded because of an error in its spike-level series.

The conditions cover an untrimmed Dorado baseline (C0 and C1), Dorado demultiplexing with integrated trimming (C2 and C3), Barbell 0.3.2 in its default kit mode applied directly to the untrimmed Dorado output (C4 and C5; S5.2), and a two-stage Dorado-then-Barbell pipeline (C6). Basecaller release and model are confounded: HAC v6.0.0 uses the model architecture introduced with Dorado 2.0.0, whereas the Dorado 1.1.1 arm used HAC v5.0.0. The effects of release and model therefore cannot be separated.

##### S5.1 Runtime options

The following runtime options were set for the tools indicated:

- `--threads` 56 or 64 (Barbell); `-@ 120` (samtools)
- `--device auto` (Dorado GPU/CPU auto-detection)
- `--recursive` (Dorado: recurse into POD5 subdirectories)

##### S5.2 Barbell invocation (C4, C5, C6)

Barbell 0.3.2 was run in kit mode (`barbell kit --kit SQK-NBD114-96`) with its default filter pattern set. The logs of all 24 Barbell runs (C4–C6 of eight experiments) report the kit type “Safe”. Search thresholds and the `--use-extended` option were left at their defaults. In the released driver, this setting corresponds to `barbell_paper.py --barbell-mode default`. The driver’s own default, `--barbell-mode maximize`, adds the `--maximize` flag and was not used. The reported results were produced with the following command:

```
barbell kit --kit SQK-NBD114-96 --threads {threads} \
  --input {input_fastq} \
  --output {output_dir}
```

For C6, the driver concatenated all `*.fastq` files in the C3 output directory and its subdirectories, including Dorado’s unclassified reads, into `merged_c3.fastq` and ran Barbell with the same parameters as for C4 and C5.

The default pattern set accepts a single barcode at either read end or concordant barcodes at both ends. Barbell writes one trimmed FASTQ file per barcode (`NBxx.trimmed.fastq`) and no file for unclassified reads. C4–C6 therefore contain only reads with an accepted barcode configuration, whereas the Dorado outputs C2 and C3 include Dorado’s unclassified reads.

##### S5.3 BAM → FASTQ conversion

The BAM files written by Dorado 2.0.0 (C1 and the per-barcode C3 files) were converted to FASTQ with `samtools fastq`; the option `-T "*"` copies all BAM auxiliary tags into the FASTQ header comments:

```
samtools fastq -T "*" -@ 120 {input.bam} > {output.fastq}
```

#### S6. Residual barcode scanning

Residual barcodes were detected by a fuzzy regular-expression search for the 192 barcode orientations in every FASTQ file of a condition’s output directory, so the C2 and C3 populations include Dorado’s unclassified output.

#### S6.1 Fuzzy matching parameters

Each of the 192 barcode orientations was compiled with the Python `regex` package as `(SEQUENCE){e<=2}` with the `IGNORECASE` and `BESTMATCH` flags (`--max-edits 2`). A match may therefore differ from the 24 nt barcode by up to two substitutions, insertions, or deletions in any combination (Levenshtein distance ≤ 2), so a matched span is 22–26 nt long.

The detector (`compare_dorado_barbell_outputs.py --detector occurrences`, which uses `OccurrenceScanner` in `validate_detector_occurrences.py`) reports every non-overlapping occurrence in a read, including repeated copies of the same barcode orientation. Each 24 nt barcode orientation is split into three non-overlapping 8 nt seeds that are matched exactly. Two edits can alter at most two of the three seeds, so every match within the edit budget contains at least one intact seed. Each seed hit defines candidate starts, shifted by up to 2 nt in either direction to allow for insertions or deletions upstream of the seed. Each candidate span of 22–26 nt is then confirmed by a full match of the same pattern, and `BESTMATCH` keeps its reported edit count minimal. Overlapping candidates from all 192 patterns are resolved greedily in the order edit distance, deviation from 24 nt, start, end, barcode ID, and orientation: a candidate that overlaps an already retained occurrence by at least 1 nt is discarded, whereas adjacent occurrences are kept. `residual_count` is the number of retained occurrences. Each output directory records the detector, the SHA-256 checksum of the scanner source, the regex version, the edit budget, and the terminal window.

#### S6.2 Position classification

Match positions were classified with `--terminal-window 80`. With zero-based, end-exclusive coordinates, a match is terminal if `start` ≤ 80 or `read_length` − `end` ≤ 80, and internal otherwise. Per-read flags were:

- `terminal`: at least one match within 80 nt of either the 5′ or the 3′ read end.
- `internal`: at least one match located entirely outside both 80 nt terminal windows.
- `both_ends`: terminal matches detected at both the 5′ and the 3′ end of the same read.

The flags use all retained occurrences, so a read with both a terminal and an internal match counts toward both totals. In short reads, the two end windows overlap: a single 24 nt match at positions 20–44 of a 64 nt read passes both end-window tests, so `both_ends` does not necessarily mean two occurrences.

##### S6.3 Pattern classes

In addition to these flags, each read was assigned exactly one pattern class (`residual_pattern_summary.csv`):

- `no_residual_barcode`
- `single_residual_barcode__{terminal | internal | both_ends}`
- `multiple_same_barcode_same_direction__{terminal | internal | both_ends}`
- `multiple_same_barcode_mixed_direction__{terminal | internal | both_ends}`
- `multiple_different_barcodes__{terminal | internal | both_ends}`

The location suffix follows the priority `internal > both_ends > terminal`: a read with any internal match is `__internal`, a read with terminal matches at both ends but no internal match is `__both_ends`, and all other positive reads are `__terminal`. Unlike the flags of S6.2, the pattern classes, and therefore the location composition in Figure 1G, are mutually exclusive. "Single" means one detected occurrence, "`same_direction`" repeated copies of one barcode orientation, and "`mixed_direction`" matches to both orientations of one barcode ID. `single_residual_barcode__both_ends` is possible only in short reads, in which the two end windows overlap (S6.2).

Counting occurrences rather than distinct patterns, a single copy was found in 90.3% (C0), 89.1% (C1), 70.8% (C2), 71.3% (C3), 99.4% (C4 and C5), and 98.8% (C6) of positive reads, and three or more copies in 5.7% (C2) and 6.1% (C3). Reads with repeated copies of one orientation (`multiple_same_barcode_same_direction`) occurred at 287 and 310 reads per million output reads (RPM; pooled) in C2 and C3, 23 and 28 RPM in C4 and C5, and 139 RPM in C6.

Counted as distinct barcode-orientation patterns (Figure 1D), one pattern was found in 90.6% (C0), 89.3% (C1), 72.5% (C2), 72.9% (C3), and 99.6% (C4 and C5) of positive reads. C2 and C3 had two patterns in 23.4% and 22.7% and three or more in 4.1% and 4.5% of positive reads. Reads matching more than one distinct barcode ID (Figure 1E) occurred at 4,050 and 4,571 RPM in C0 and C1, at 3,322 and 3,828 RPM in C2 and C3, and at 3.3 and 4.9 RPM in C4 and C5 (geometric means of `multiple_barcode_id_reads` per million output reads; `per_experiment_results.csv`). Barcode-positive reads fell from 828,550 to 14,173 RPM between C0 and C2 and from 850,109 to 16,590 RPM between C1 and C3. The multi-ID share of positive reads therefore rose after Dorado demultiplexing, although the rate of multi-ID reads changed little.

##### S6.4 NanoStat QC

Per-condition read-length and Q-score statistics were generated with NanoStat 1.6.0. Read Q-scores are the per-read quality values predicted by the basecaller. NanoStat reports their median and a mean that converts each read's Q-score to an error probability, averages these probabilities, and converts the average back to a Q-score. Because low-quality reads dominate the averaged error probability, this mean is lower than the median. For the rebasecalled C0 output, NanoStat reports per-experiment median read Q-scores of 14.1–15.3 and mean read Q-scores of 11.3–13.6. The corresponding C1 ranges are 13.2–13.9 and 10.8–12.6. The MinKNOW sequencing summaries of the live basecalling give higher values for the pass reads (median 16.6–18.3 per experiment; 14.8–16.3 with the NanoStat calculation of the mean; `adaptive_sampling_pass_reads.csv`). These values are not directly comparable: the rebasecalled output was produced with other HAC models (v5.0.0 and v6.0.0 instead of v4.3.0) and also contains the fail reads (S3), whereas MinKNOW pass reads have a mean Q-score of at least 9. None of these values measures empirical read accuracy.

NanoStat was run per condition with `--name {condition}`; its text summaries are stored as `{condition_dir}/nanostat/{condition}` (e.g., `exp_c0_dorado111_raw/nanostat/c0`).

##### S6.5 Database backend and parallel processing

Read-level results (read length, Q-score statistics, residual counts, and pattern class) were stored in a per-experiment SQLite database (`comparison.sqlite`). Scanning ran in parallel with 56 worker processes (Python `concurrent.futures.ProcessPoolExecutor`), with 5,000 reads per work item and database inserts in batches of 500,000 reads.

All seven conditions of an experiment were scanned together with the following command, run from the project root with the barcode file of S2 (`barcodes.csv`). `{decoys}` was 2 for Exp 4, Exp 6, and Exp 7 (S6.7) and 0 otherwise. The options `--skip-nanostat` and `--skip-barcode-scan` skip the NanoStat step (run separately; S6.4) and the script's optional minimap2 barcode scan. They do not affect the regex detector:

```
python3 scripts/compare_dorado_barbell_outputs.py \
  --condition-dirs {dir_c0} {dir_c1} {dir_c2} {dir_c3} {dir_c4} {dir_c5} {dir_c6} \
  --condition-names c0 c1 c2 c3 c4 c5 c6 \
  --barcode-csv barcodes.csv --out {experiment_output_dir}/cmp_all \
  --detector occurrences --max-edits 2 --decoys {decoys} --decoy-seed 20260929 \
  --threads {threads} --skip-nanostat --skip-barcode-scan --overwrite
```

##### S6.6 Synthetic controls

Unit tests checked the occurrence scanner with the edit budget of S6.1 (`code/scripts/tests/test_detector_occurrences.py`). In 60 random trials, a 24 nt test barcode carrying zero to three random edits was placed between random flanks. In every trial, the seed-based candidates equaled the set of all 22–26 nt spans with an edit distance of at most 2 found by exhaustive dynamic programming. Every variant with two deletions or two insertions, at every pair of positions, was detected as exactly one match, and a variant with three substitutions was rejected.

The decoy scan of the production script (S6.7) was checked by planting each sequence of both decoy sets in the middle of a random read with 300 nt flanks, either unchanged, with one substitution, insertion, or deletion, or with each combination of two of these edits. All 3,840 planted copies (2 sets × 192 decoys × 10 variants) were counted as internal decoy matches (`code/scripts/tests/test_decoy_planted_copies.py`).

##### S6.7 Decoy background

To estimate the chance-match background of the detector, all 62,550,076 output reads of the three smallest experiments (Exp 4, Exp 6, and Exp 7; all seven conditions) were also scanned against two decoy barcode sets (`compare_dorado_barbell_outputs.py --decoys 2`, seed 20260929). Each decoy set shuffles the bases of every forward barcode, which preserves base composition, and uses the reverse complement of the shuffled sequence as the backward decoy, so the forward/reverse-complement pairing of the real set is kept. Decoy matches were located and classified exactly like real matches (Table S3). The C6 outputs of these experiments (17,036–47,465 reads) contained no decoy match. Terminal decoy rates were at most 1.1 RPM, and no decoy read was classified as both-ends.

#### S7. Cross-experiment aggregation and normalization

Cross-experiment summary statistics were produced by `barbell_figures_summa.py` from the eight per-experiment `cmp_all/` and `figures/` directories. The default summary is the mean ± standard deviation (SD) across experiments; the standard error (option `--use-se`) was not used. Read retention is reported pooled, as the sum of retained reads divided by the sum of input reads across experiments. Pooled values differ by up to 1.2 percentage points from the mean of the per-experiment ratios (largest difference: C4, 91.8% pooled versus 90.6%). S7.3 lists the summaries used in each figure.

##### S7.1 Reads-per-million normalization

Library sizes ranged from about 1.75 to 15.4 million input reads, so absolute residual counts mainly reflect library size. Read-location counts (terminal, internal, and both-ends) and per-barcode-ID residual counts were therefore normalized per million output reads of the same condition and experiment before averaging.

##### S7.2 Summary outputs and Supplementary Data 1

`barbell_figures_summa.py` writes wide summary CSVs (`condition`, `group`, `stat_n`, `stat_mean`, `stat_std`, `stat_se`, `stat_min`, and `stat_max`, followed by one column per experiment) and long-format per-experiment tables. Supplementary Data 1 (`supplementary_source_data.zip`) contains the per-experiment results underlying the figures and tables. It contains no identifying donor-level data, read-level data, read identifiers, input sequences, acquisition/QC reports, or local paths. Besides `README.md`, which describes every file, the archive contains:

- `read_level/`: `per_experiment_results.csv` (one row per experiment and condition: read counts, median read length, barcode-positive, internal, and multiple-barcode-ID reads and percentages, location classes per million output reads, pattern and occurrence compositions, and reads in Dorado's unclassified output for C2 and C3; Figures 1B–1G and S1A, Results, and S10.1), `condition_summary.csv` (mean, SD, median, quartiles, minimum, maximum, geometric mean, geometric standard deviation (GSD), and pooled value per condition; Table S4), `paired_internal_comparisons.csv` (Table S6), `decoy_background_per_experiment.csv` (Table S3), `figureS1_read_length_counts_10bp.csv` (pooled read counts per 10 nt bin; Figure S1B–H), and `adaptive_sampling_pass_reads.csv` (adaptive-sampling rejections and read Q-scores of the pass reads in the MinKNOW sequencing summaries; S3 and S6.4).
- `figure2/`: `audit_summary.json` (Table S7), `blast_provenance.json`, `construct_context_pairs.csv` (barcode, TaxID group, record category, and longest construct per barcode-accession pair, without accessions), `construct_crosscheck_summary.json`, `Fig2A_taxid_groups.csv` and `Fig2A_taxid_groups_named.csv` (Figure 2A), `Fig2B_barcode_ranking.csv`, `Prism_Fig2B_longest_construct.csv` (Figure 2B, rendered in R), `Fig2B_single_accession_barcodes.csv` (Table S8), `Fig2C_composition.csv`, `Prism_Fig2C_B24_pair_counts.csv` (Figure 2C), and `taxonomy_mapping_summary.json`.

- `code/scripts/`: the scripts used for the analyses and figures (section S4) and, in `code/scripts/tests`, the two unit tests cited in S6.6.

##### S7.3 Summary figure conventions

Figure 1B shows the mean of the eight per-experiment medians, and Figure 1D the mean per-experiment compositions. Figures 1E–1G show geometric means with the GSD. Unless stated otherwise, all other summaries are arithmetic means ± SD. The rates in Figures 1E–1G differ between conditions and location classes by more than four orders of magnitude and vary up to almost 90-fold between the eight experiments of one condition (Table S4). The GSD is the corresponding multiplicative dispersion factor, so geometric mean ×/÷ GSD is the log-scale analog of mean ± SD.

Figures S1B–H use pooled read counts: the summed read count of a 10 nt length bin across the eight experiments is divided by the largest such bin of the same condition (peak = 1). Median read lengths and the read-length histograms were computed from the per-read lengths recorded in the residual-scan database (S6.5).

##### S7.4 Residual barcode location composition (Figure 1G)

The location composition in Figure 1G was derived from the per-experiment files `cmp_all/tool_summary.csv` (total reads) and `cmp_all/residual_pattern_summary.csv` (pattern class of each read). For each condition and each of the three mutually exclusive location classes (terminal at one end only, internal, and both-ends; S6.3), the per-experiment read count was normalized per million output reads. Table S4 summarizes the eight per-experiment values. In Figure 1G, each symbol is one experiment, the horizontal line marks the geometric mean, and the error bars span geometric mean ×/÷ GSD (S7.3). The rates span more than four orders of magnitude, so the y-axis is logarithmic.

#### S8. Statistical analysis

Statistical comparisons use paired experiment-level internal-match rates (S8.2). Read-level binary outcomes and length distributions are summarized descriptively.

##### S8.1 Descriptive binary outcomes

The binary outcomes in Table S5 define descriptive per-read measurements. Conditions share source reads, so their read-level counts are not treated as independent samples for significance testing.

##### S8.2 Paired cross-experiment analysis of internal residuals

To test whether the reduction of internal residual barcodes was consistent across the eight experiments, each experiment was treated as a paired observation, and the internal residual rate (RPM) was compared between conditions with the two-sided Wilcoxon signed-rank test (`scipy.stats.wilcoxon`). Table S6 gives the per-experiment fold-changes as median (range). The main text reports the geometric mean of the same ratios, which equals the ratio of the geometric means in Table S4. With n = 8, the smallest attainable two-sided p-value is $2/2^8 = 0.0078$, reached only when all eight experiments change in the same direction.

In all eight experiments, detected internal rates were lower after Barbell than after Dorado demultiplexing within the same basecaller lineage: median 48.3-fold for C2 versus C4 and 52.0-fold for C3 versus C5 (geometric means 50.0 and 50.7). Dorado-demultiplexed output had about 2.2-fold lower detected internal rates than its untrimmed baseline (C0 versus C2 and C1 versus C3), and C3 had higher detected internal rates than C2 in all eight experiments.

#### S9. Reference database screen for ONT barcodes

The 96 SQK-NBD114.96 barcode sequences (24 nt each) were queried alone and within three native constructs (S9.1) against the NCBI core_nt database with BLASTn (BLAST+ 2.15.0+). The core_nt release was dated 12 September 2026 and contained 130,787,235 sequences totaling 1,079,005,865,156 bases (longest sequence 98,200,793 bases) in 91 volumes; `blastdbcheck` reported no errors.

##### S9.1 Query construction and search conditions

Four nested query panels were searched, each with 192 queries (96 barcodes in two orientations). Construct sequences were taken from Dorado 1.1.1, the release used for C0 and C2: the barcode flanks of the `kit_nb96` definition (`barcode_kits.cpp`) and the adapter sequence that Dorado assigns to this kit, the LSK110 front adapter (`adapter_primer_kits.cpp`). Panel names give the content (B, barcode only; K, kit construct) and the query length in nt. Because each construct must match over its full length (S9.3), a construct match extends the identity from the barcode into the adjoining kit sequence. The panels were built with `make_construct_panels.py`:

- B24 (barcode only, 24 nt): each barcode in forward and reverse-complement orientation, as in the residual scan.

- K40 (ligation flanks, 40 nt): `AAGGTTAA` + barcode + `CAGCACCT`, the barcode with eight bases of native flank on each side.
- K46 (Dorado window, 46 nt): `ATTGCTAAGGTTAA` + barcode + `CAGCACCT`, the complete front and rear flanks that Dorado searches together with the barcode for this kit.
- K68 (adapter construct, 68 nt): the LSK110 front adapter `CCTGTACTTCGTTCAGTTACGTATTGC` joined to the K46 window. The adapter ends and the front flank begins with `ATTGC`, which is included once, giving `CCTGTACTTCGTTCAGTTACGTATTGCTAAGGTTAA` + barcode + `CAGCACCT`.

For K40, K46, and K68, the second query of each barcode is the reverse complement of the whole construct, that is, the sequence expected at the 3′ end of a read. For B24 it is the reverse-complement barcode. Because BLASTn searched both strands, the second orientation is redundant and serves as a symmetry check. The counts in S9.6 account for this redundancy. The barcode occupies query positions 1–24 (B24), 9–32 (K40), 15–38 (K46), and 37–60 (K68) in the forward constructs and 9–32 in the reverse-complement constructs. All 192 queries of a panel are distinct. The kit's barcode set contains no palindromic or duplicated barcodes. In total, 768 queries were built.

In the first 200,000 reads each of C0 and C1 from two experiments (Exp 1 and Exp 6; 800,000 reads), exact barcode matches were located in the first and last 150 nt, and base frequencies were tabulated per position for 40 nt upstream and 12 nt downstream of the barcode. The majority sequence was `TTCAGTTA CGTATTGCTAAGGTTAA` + barcode + `CAGCACCT` at read starts and `AGGTGCTG` + reverse-complement barcode + `TTAACCTTAGCAAT` at read ends, matching the K68 construct and the reverse complement of the K46 window. Of the reads with an exact barcode in the first 150 nt, 91.9–93.4% carried the complete K40 construct and 79.5–84.1% the complete K46 window, but only 2.5–7.3% the complete K68 construct, because the first adapter bases at read starts were often missing or miscalled.

##### S9.2 BLASTn parameters and database

BLASTn from BLAST+ 2.15.0 was run against core_nt with `-task blastn-short`, `-evalue 10`, `-dust no`, `-soft_masking false`, and `-max_target_seqs 5000` for all four panels (`run_barcode_core_nt_blast.sh`, 64 threads). The e-value threshold was set explicitly because the `blastn-short` default differs between BLAST+ versions. Word size, scoring, and strand were left at their defaults, so both strands were searched. The tabular output (`-outfmt 6`) contained the fields `qseqid`, `saccver`, `staxids`, `sscinames`, `pident`, `length`, `qlen`, `qcovs`, `mismatch`, `gapopen`, `qstart`, `qend`, `sstart`, `send`, `evalue`, `bitscore`, and `stitle`. These permissive settings also return partial alignments, so the strict criterion of S9.3 was applied afterward to every exported row. The raw output comprised 3,210 (B24),

4,110 (K40), 4,264 (K46), and 122,288 (K68) high-scoring segment pairs (HSPs), and 174, 186, 188, and 192 of the 192 queries per panel returned at least one alignment. The `-max_target_seqs` ceiling was not approached: the largest number of distinct subject sequences for any single query was 194 (B24), 582 (K40), 583 (K46), and 1,077 (K68), below the predefined stop threshold of 4,500.

##### S9.3 Strict complete-query identity audit

The strict post-hoc audit requires `pident` = 100%, `mismatch` = 0, `gapopen` = 0, `min(qstart,qend)` = 1, `max(qstart,qend)` = `qlen`, `length` = `qlen`, and `abs(send - sstart)` + 1 = `qlen`, with positive query length and subject coordinates. The alignment thus covers the complete query, in either coordinate orientation, with an ungapped subject span of the same length. The output has no separate `gaps` column, but zero gap openings together with equal alignment, query, and subject lengths exclude gaps. The rule retains 1,390 (B24), 574 (K40), 424 (K46), and six (K68) HSPs. The `qcovs` field cannot replace these checks, because it aggregates query coverage across all HSPs of a subject: among rows with `pident` = 100%, `mismatch` = 0, and `qcovs` = 100%, the rule rejects 102 (B24), 148 (K40), 166 (K46), and four (K68) partial alignments, all of which fail the complete query-span, alignment-length, and subject-span checks.

##### S9.4 Genomic-element classification

Subject titles (`stitle`) were classified by keyword rules applied in a fixed order: plasmid; rRNA gene (16S, 18S, ribosomal RNA, or rRNA); phage/virus/viral; organelle (mitochondrial, chloroplast, plastid, or apicoplast terms); chromosome/genome (chromosome, complete genome, or genome assembly); and other/uncultured. The first matching rule supplies the label. The chromosome/genome label is deliberately neutral, because a title does not establish bacterial origin, and taxonomy grouping (S9.5) is independent of these rules. The primary composition counts each of the 505 distinct barcode-accession pairs once, collapsing repeated HSPs and both query orientations; the HSP-weighted composition (n = 1,390) is secondary.

Sixty-four pairs (12.7%) and 198 HSPs (14.2%) match keywords of more than one category: chromosome/genome with phage/virus (53 pairs), plasmid (four), or organelle (six), and phage/virus with plasmid (one); the corresponding HSP counts are 156, eight, 32, and two. No retained title matches both plasmid and rRNA keywords. Moving chromosome/genome to the first position would change its pair count from 117 to 180 (23.2% to 35.6%), phage/virus from 63 to ten, organelle from 19 to 13, and plasmid from 113 to

109 (HSPs: 374 to 570, 180 to 24, 60 to 28, and 264 to 256); rRNA gene and other counts would not change. The specific labels were kept ahead of the generic genome rule, so that the generic label applies only to records without a more specific keyword.

Across the 505 barcode-only pairs, rRNA gene, chromosome/genome, plasmid, phage/virus, organelle, and other/uncultured records contributed 180 (35.6%), 117 (23.2%), 113 (22.4%), 63 (12.5%), 19 (3.8%), and 13 (2.6%) pairs, respectively (81.2% for the first three combined). Weighted by HSPs, the shares were 34.5%, 26.9%, 19.0%, 12.9%, 4.3%, and 2.3%. The same rules classify the construct panels (`Fig2C_composition.csv`). Among the 224 K40 pairs, chromosome/genome, plasmid, rRNA gene, phage/virus, organelle, and other/uncultured records contributed 28.6%, 23.7%, 19.6%, 17.9%, 7.1%, and 3.1%. Among the 170 K46 pairs, 35.9%, 25.3%, 14.7%, 14.1%, 7.1%, and 2.9%.

##### S9.5 Per-barcode metrics

Per-barcode and per-group summaries use the strict retained set. Taxonomy grouping uses the exported `staxids` field (NCBI Taxonomy identifier, TaxID). Each of the 1,390 retained barcode-only rows contains one TaxID, giving 195 distinct TaxID groups. Names were obtained from NCBI Taxonomy EFetch (`https://eutils.ncbi.nlm.nih.gov/entrez/eutils/efetch.fcgi`; `db=taxonomy`, `retmode=xml`). The lookup on 9 October 2026 resolved all 195 TaxIDs, with no merged or missing identifiers, and returned `ScientificName`, `Rank`, `ParentTaxId`, and `LineageEx` for each (169 species, six subspecies, four strains, and 16 entries without rank; 87 bacterial, 62 eukaryotic, and 46 viral lineages). TaxID groups and numerical columns are used as exported, without regrouping by name, ancestor, or species. The species-ancestor fields of the mapping table are informational only. `Fig2A_taxid_groups_named.csv` adds the names to the plot data, and `taxonomy_mapping_summary.json` records the request URL, retrieval time, and SHA-256 checksum of the XML response. Three per-barcode counts are used, with the accession count as primary:

- primary per-barcode count: number of distinct subject accessions, equivalent to distinct barcode-accession pairs when summed across barcodes
- exported TaxID-group breadth: number of distinct `staxids` groups matched by a barcode; Figure 2A instead displays distinct barcode IDs and distinct accessions within each group
- secondary HSP count: retained alignment rows, which count both submitted orientations and every matched locus of a record. Figure 2 shows these counts descriptively, without statistical testing

Of the 192 barcode-only queries, 128 retain complete HSPs (64 barcodes in both orientations). The barcode-only set comprises 1,390 HSPs, 695 barcode/accession/coordinate loci, 505 barcode-accession pairs, and 482 distinct accessions (Table S7). NB08 has 84 accessions, 13 TaxID groups, and 184 HSPs. NB03 has 37 accessions, 22 groups, and 100 HSPs. NB04 has 35 and NB07 32 accessions. Of the 64 hit-bearing barcodes, 41 (64.1%) match more than one exported TaxID group, and 47 TaxID groups are matched by more than one barcode. NB09 has the largest TaxID-group breadth (26; 32 accessions, 90 HSPs), followed by NB03 and NB04 (22 each) and NB11 (20). *Escherichia coli* (TaxID 562; species) is matched by 17 barcode IDs, *Klebsiella pneumoniae* (TaxID 573; species) by 13, and *Acinetobacter baumannii* (TaxID 470; species) and *Cutibacterium* sp. (TaxID 1912221; NCBI rank species) by 11 each. All 96 barcodes are retained in the source ranking tables, including zero-count entries. Among TaxID groups, the largest pair count belongs to *Plasmodium berghei* ANKA (TaxID 5823; strain; Eukaryota), with 69 accessions/pairs and 146 HSPs, followed by uncultured *Muribaculaceae* bacterium (TaxID 2301481; species; Bacteria), with 31 pairs. Figure 2A retains these non-collapsed groups and labels the four highest pair counts.

##### S9.6 Count reconciliation and construct checks

Table S7 reconciles strict complete-query HSPs with loci, barcode-accession pairs, accessions, barcodes, and TaxID groups in the four panels. In every panel, the forward and reverse-complement queries returned identical sets of barcode/accession/coordinate loci, each orientation contributing one HSP per locus. The panels are nested: every K68 locus is contained in a K46 locus, every K46 locus in a K40 locus, and every K40 locus in a B24 locus of the same record. As an independent check, windows of 40 nt on either side of all 695 barcode-only loci (482 accessions) were retrieved from NCBI EFetch by versioned accession (`crosscheck_construct_loci.py`). The barcode was confirmed at every locus, and the presence of the complete K40, K46, and K68 constructs in these windows agreed with the BLAST calls at every locus (287, 212, and three positive loci, respectively), so the construct searches missed no construct locus.

The longest construct matched at each of the 505 B24 pairs was the barcode alone for 281 pairs (55.6%), the ligation flanks (K40) for 54 (10.7%), the Dorado window (K46) for 167 (33.1%), and the adapter construct (K68) for three. This classification colors the bars of Figure 2B (`Prism_Fig2B_longest_construct.csv`). Pairs carrying at least the K40 construct were frequent among organelle (16 of 19), phage/virus (40 of 63), chromosome/genome (64 of 117), and plasmid (53 of 113) records, less frequent among rRNA gene records (44 of 180), and present in seven of the 13 other/uncultured pairs. NB08 accounts for 59 of the 224 K40 pairs and 52 of the 170 K46 pairs. The three K68 pairs are a plasmid record for NB04 and a chromosome/genome and a plasmid record for NB09. A single differing flank or adapter base leaves a pair in a shorter class.

Figure 2B shows the 24 barcodes with more than three barcode-only accessions (434 pairs). The 21 barcodes with two or three accessions (52 pairs) are named in the legend of Figure 2. The remaining 19 hit-bearing barcodes match one accession each and are listed in Table S8: ten as barcode only and nine within the complete Dorado window (K46). Their records are annotated as nine plasmid, four rRNA gene, three chromosome/genome, two organelle, and one phage/virus record and belong to bacterial (12), eukaryotic (six), and viral (one) TaxIDs. Three of these records carry the barcode at two loci.

### Results

#### S10. Additional results

##### S10.1 Read yield and read-length distributions

Dorado demultiplexing retained 99.94% of the untrimmed reads in C2 and C3 (pooled counts). Dorado's unclassified output comprised 6.0% (C2) and 5.9% (C3) of the retained reads (4.7–10.6% and 4.8–10.1% per experiment). Barbell retained 91.8% (C4) and 91.2% (C5) of the reads. These output-to-input ratios do not show whether the retained reads were assigned to the correct barcode. In C6, Barbell returned 518,947 of 70,223,900 C3 reads (0.74%; mean 64,868 ± 32,614 output reads per experiment). C4 and C5 are parallel alternatives to C2 and C3, whereas C6 is an additional processing step on C3.

In the pooled read-length histograms (Figure S1B–H), bins reaching at least half of the modal height spanned 370–529 nt in C0 and 370–539 nt in C1. After Dorado demultiplexing (C2 and C3), the distribution was broader, with a shoulder toward shorter reads: the half-maximum range extended from 170 to 449 nt, and a separate minor peak at 40–49 nt reached about 20% of the modal height. After Barbell (C4 and C5), the half-maximum range (300–459 nt) was about as narrow as in untrimmed output, and the 40–49 nt peak reached 3–4% of the modal height. C6 had a pronounced 40–49 nt peak at about 40% of the modal height.

##### S10.2 Between-experiment variability

Terminal residuals in untrimmed reads were very similar across experiments (GSD 1.07 in C0 and C1; per-experiment values typically within 7% of the geometric mean). Both-ends residuals varied several-fold under every condition (GSD 1.76 in C0, rising to 4.22 in C4). In C0, they ranged from 21,365 to 145,634 RPM, an approximately 7-fold range (median 78,685 RPM; IQR 64,893–98,423). The experiments kept a consistent order across conditions: Exp 7 had the lowest internal rate under all seven conditions and Exp 6 the highest under six of seven. For both-ends residuals, Exp 7 was lowest in six and Exp 6 highest in all seven conditions. A large part of the spread between experiments is therefore experiment-specific rather than tool-dependent, and the paired analysis (S8.2) removes this part. The order was reversed only for terminal residuals in untrimmed output (C0 and C1). There, terminal matches account for about 75% of all reads, and the class priority (internal > both-ends > terminal) moves reads with an additional internal or both-ends match out of the terminal class, so experiments with high internal and both-ends rates have lower terminal rates.

Each experiment comprises the six samples (T0–T5) of one donor, whose libraries were prepared from 13.2–25 µL of DNA extract (S2). Donor, DNA input, and sequencing run therefore cannot be separated as causes. Exp 6 and Exp 7 were also the two smallest experiments (1.75 and 3.75 million reads in C0, compared with 5.3–15.4 million in the other six experiments). Because they lie at opposite ends of the ranking, library size alone does not explain the order, and the correlation of library size with the internal rate was weak (Spearman rho −0.24 to −0.40 across conditions; with the both-ends rate, −0.21 to −0.52).

#### Abbreviations

| Abbreviation | Full term |
|---|---|
| **BAM** | Binary Alignment/Map format |
| **BLAST/BLASTn** | (nucleotide) Basic Local Alignment Search Tool |
| **CPU/GPU** | central/graphics processing unit |
| **CSV** | comma-separated values (file format) |
| **DNA** | deoxyribonucleic acid |
| **EDTA** | ethylenediaminetetraacetic acid |
| **FAST5 / POD5** | ONT raw-signal file formats |
| **FASTQ** | text-based sequencing-read format containing nucleotide sequences and quality scores |
| **GSD** | geometric standard deviation |
| **HAC** | high-accuracy basecalling model |
| **hg38** | human reference genome assembly GRCh38 |
| **HSP** | high-scoring segment pair (BLAST alignment) |
| **ID** | identifier |
| **IQR** | interquartile range |
| **JSON** | JavaScript Object Notation (file format) |
| **LSK110** | Ligation Sequencing Kit 110 (Oxford Nanopore Technologies) |
| **NB01–NB96** | native barcodes 1–96 of the SQK-NBD114.96 kit |
| **NCBI** | National Center for Biotechnology Information |
| **nt** | nucleotides |
| **ONT** | Oxford Nanopore Technologies |
| **Q1/Q3** | first/third quartile |
| **Q-score** | Phred quality score |
| **QC** | quality control |
| **RPM** | reads per million |
| **rRNA** | ribosomal ribonucleic acid |
| **SD** | standard deviation |
| **SE** | standard error |
| **SHA-256** | Secure Hash Algorithm 256-bit |
| **SQK-LSK114** | Ligation Sequencing Kit V14 (Oxford Nanopore Technologies) |
| **SQK-NBD114.96** | Native Barcoding Kit 96 V14 (Oxford Nanopore Technologies) |
| **TaxID** | NCBI Taxonomy identifier |
| **XML** | Extensible Markup Language |

**Table S1.** Software versions used in this study.

| Software | Version | Notes / invocation |
|---|---|---|
| Dorado | 1.1.1+e72f1492 | Direct command-line call (system `PATH`) |
| Dorado | 2.0.0+20e87c8b | Run in a conda environment (`conda run --no-capture-output -n dorado200 dorado`) |
| Barbell | 0.3.2 | Bioconda package (build `h54198d6_0`) in a separate conda environment |
| NanoStat | 1.6.0 | Per-condition read-length and Q-score QC |
| SAMtools | 1.19.2 | BAM-to-FASTQ conversion (`samtools fastq`) for C1 and the per-barcode C3 BAM files |
| Python | 3.12 | Analysis scripts, including the BLAST audit and taxonomy mapping (section S9); packages `regex` 2026.4.4, `numpy`, `scipy`, and `matplotlib` |
| BLAST+ (blastn) | 2.15.0+ | BLASTn search followed by a strict complete-query, ungapped identity audit (section S9) |
| R | 4.5.3 (2026-03-11) | Rendering of Figures 2A and 2B (`plot_audited_figure2_graphpad.R`; packages `ggplot2` and `readr`) |
| pod5 | 0.3.39 | FAST5-to-POD5 conversion (command `pod5 convert fast5`) |
| GraphPad Prism | 11 | Rendering of Figures 1, 2C, and S1 from the exported tables |

**Table S2.** Seven-condition preprocessing matrix.

| ID | Basecaller | HAC model | Demultiplexer | Trimming | Input → output |
|---|---|---|---|---|---|
| C0 | Dorado 1.1.1 | v5.0.0 | none | `--no-trim` | POD5 → `exp_c0_dorado111_raw/merged_raw.fastq` |
| C1 | Dorado 2.0.0 | v6.0.0 | none | `--no-trim` | POD5 → `exp_c1_dorado200_raw/merged_raw.fastq` |
| C2 | Dorado 1.1.1 | v5.0.0 | Dorado | default (on) | POD5 → `exp_c2_dorado111_demux/` (per-barcode FASTQ, `--emit-fastq`) |
| C3 | Dorado 2.0.0 | v6.0.0 | Dorado | default (on) | POD5 → `exp_c3_dorado200_demux/` (per-barcode BAM → FASTQ) |
| C4 | Dorado 1.1.1 | v5.0.0 | Barbell 0.3.2 | Barbell-internal | C0 `merged_raw.fastq` → `exp_c4_barbell111_default/barbell_demux/` |
| C5 | Dorado 2.0.0 | v6.0.0 | Barbell 0.3.2 | Barbell-internal | C1 `merged_raw.fastq` → `exp_c5_barbell200_default/barbell_demux/` |
| C6 | Dorado 2.0.0 | v6.0.0 | Dorado → Barbell | two-stage | All C3 FASTQ files, including unclassified, concatenated → `exp_c6_dorado200_barbell_default/barbell_demux/` |

BAM, Binary Alignment/Map; FASTQ, text-based read format with quality scores; HAC, high-accuracy basecalling model; POD5, ONT raw-signal file format.

**Table S3.** Decoy background in Exp 4, Exp 6, and Exp 7 (reads per million output reads of each condition; decoy reads in parentheses).

| Experiment | Condition | Reads | Decoy 1 internal RPM | Decoy 2 internal RPM | Real internal RPM | Decoy terminal RPM |
|---|---|---|---|---|---|---|
| Exp 4 | C0 | 5,326,744 | 2.3 (12) | 0.8 (4) | 6,931 | 0.6 / 0.4 |
| | C1 | 5,327,218 | 2.3 (12) | 1.1 (6) | 7,937 | 0.8 / 0.4 |
| | C2 | 5,322,542 | 2.1 (11) | 0.2 (1) | 2,736 | 0.4 / 0.8 |
| | C3 | 5,322,841 | 1.9 (10) | 0.4 (2) | 2,997 | 0.4 / 0.8 |
| | C4 | 4,701,618 | 1.9 (9) | 0.6 (3) | 37.2 | 0.4 / 0.2 |
| | C5 | 4,716,641 | 1.9 (9) | 0.6 (3) | 42.8 | 0.4 / 0.6 |
| | C6 | 47,465 | 0.0 (0) | 0.0 (0) | 337 | 0.0 / 0.0 |
| Exp 6 | C0 | 1,747,329 | 0.6 (1) | 1.1 (2) | 6,622 | 0.6 / 0.0 |
| | C1 | 1,747,179 | 1.1 (2) | 1.7 (3) | 8,261 | 0.6 / 0.0 |
| | C2 | 1,743,432 | 0.0 (0) | 0.6 (1) | 3,670 | 1.1 / 0.6 |
| | C3 | 1,743,027 | 0.6 (1) | 1.1 (2) | 4,605 | 0.6 / 0.6 |
| | C4 | 1,483,743 | 0.7 (1) | 0.0 (0) | 94.4 | 0.0 / 0.0 |
| | C5 | 1,477,781 | 1.4 (2) | 0.7 (1) | 86.6 | 0.7 / 0.0 |
| | C6 | 17,036 | 0.0 (0) | 0.0 (0) | 1,057 | 0.0 / 0.0 |
| Exp 7 | C0 | 3,751,815 | 1.1 (4) | 1.9 (7) | 1,294 | 0.0 / 0.3 |
| | C1 | 3,751,744 | 1.1 (4) | 1.6 (6) | 1,359 | 0.0 / 0.3 |
| | C2 | 3,748,321 | 0.8 (3) | 1.6 (6) | 504 | 0.3 / 0.5 |
| | C3 | 3,748,251 | 0.5 (2) | 1.3 (5) | 508 | 0.3 / 0.5 |
| | C4 | 3,408,222 | 0.9 (3) | 1.2 (4) | 15.0 | 0.0 / 0.6 |
| | C5 | 3,388,412 | 0.9 (3) | 0.9 (3) | 17.4 | 0.0 / 0.9 |
| | C6 | 28,715 | 0.0 (0) | 0.0 (0) | 69.7 | 0.0 / 0.0 |

Real internal rates are from the occurrence detector on the same reads. Terminal decoy rates are given for decoy set 1 / decoy set 2.

**Table S4.** Per-condition, per-location descriptive statistics of residual barcode burden across the eight experiments, in reads per million output reads (RPM).

| Condition | Location | n | Min | Q1 | Median | Q3 | Max | IQR | Geometric mean | GSD |
|---|---|---|---|---|---|---|---|---|---|---|
| C0 | terminal | 8 | 635,495 | 724,633 | 764,016 | 774,851 | 790,574 | 50,218 | 742,441 | 1.07 |
| | internal | 8 | 1,294 | 2,998 | 4,232 | 5,397 | 6,931 | 2,399 | 3,816 | 1.73 |
| | both_ends | 8 | 21,365 | 64,893 | 78,685 | 98,423 | 145,634 | 33,531 | 72,085 | 1.76 |
| C1 | terminal | 8 | 644,166 | 741,284 | 769,203 | 783,359 | 798,148 | 42,075 | 751,724 | 1.07 |
| | internal | 8 | 1,359 | 3,225 | 4,714 | 6,019 | 8,261 | 2,795 | 4,227 | 1.80 |
| | both_ends | 8 | 24,995 | 72,165 | 86,843 | 116,376 | 162,616 | 44,210 | 82,373 | 1.76 |
| C2 | terminal | 8 | 3,196 | 4,462 | 5,011 | 5,847 | 7,201 | 1,386 | 4,945 | 1.36 |
| | internal | 8 | 504 | 1,330 | 1,879 | 2,665 | 3,670 | 1,334 | 1,730 | 1.85 |
| | both_ends | 8 | 990 | 4,389 | 8,640 | 12,142 | 29,079 | 7,753 | 6,476 | 3.14 |
| C3 | terminal | 8 | 4,357 | 5,417 | 6,088 | 6,735 | 8,140 | 1,318 | 5,996 | 1.26 |
| | internal | 8 | 508 | 1,389 | 2,013 | 2,820 | 4,605 | 1,431 | 1,866 | 1.94 |
| | both_ends | 8 | 1,128 | 5,057 | 9,891 | 13,759 | 32,873 | 8,701 | 7,377 | 3.15 |
| C4 | terminal | 8 | 4,822 | 8,737 | 9,598 | 12,811 | 21,710 | 4,073 | 10,368 | 1.56 |
| | internal | 8 | 15.0 | 26.5 | 32.8 | 41.6 | 94.4 | 15.0 | 34.6 | 1.73 |
| | both_ends | 8 | 97.1 | 1,018 | 1,450 | 2,131 | 8,506 | 1,113 | 1,121 | 4.22 |
| C5 | terminal | 8 | 5,002 | 8,465 | 9,129 | 11,481 | 20,906 | 3,016 | 9,766 | 1.50 |
| | internal | 8 | 17.4 | 27.8 | 36.7 | 45.7 | 86.6 | 17.9 | 36.8 | 1.64 |
| | both_ends | 8 | 109 | 1,111 | 1,681 | 2,226 | 9,458 | 1,114 | 1,234 | 4.17 |
| C6 | terminal | 8 | 9,786 | 23,768 | 30,510 | 35,768 | 53,592 | 12,000 | 27,316 | 1.67 |
| | internal | 8 | 69.7 | 290 | 305 | 376 | 1,057 | 86.6 | 311 | 2.13 |
| | both_ends | 8 | 668 | 6,636 | 15,421 | 20,102 | 34,222 | 13,467 | 9,655 | 3.52 |

Q1/Q3, first/third quartile computed by linear interpolation between the closest ranks (`numpy.percentile` default); IQR, interquartile range (Q3 − Q1). Geometric mean is $10^{\mathrm{mean}(\log_{10}(x))}$ across the eight experiments; GSD is $10^{\mathrm{SD}(\log_{10}(x))}$ (S7.3). Values ≥ 100 are rounded to the nearest integer; values < 100 to one decimal.

**Table S5.** Per-read binary outcomes used for descriptive comparisons.

| Outcome | Definition |
|---|---|
| `residual_barcode_present` | `residual_count` > 0 (any fuzzy match anywhere in the read) |
| `internal_residual_barcode_present` | `internal_residual_count` > 0 (match outside both 80 nt terminal windows) |
| `multiple_different_residual_barcodes` | More than one distinct barcode ID detected in the same read |
| `both_end_residual_barcode` | `both_ends_residual` = 1 (terminal matches at both the 5′ and 3′ end) |
| `short_read_lt_250bp` | `read_length` < 250 nt |
| `short_read_lt_500bp` | `read_length` < 500 nt |

**Table S6.** Paired cross-experiment comparison of the internal residual barcode rate (reads per million output reads; n = 8 experiments).

| Comparison | Median RPM (A) | Median RPM (B) | Median fold-change | Fold-change range | Runs A>B | Wilcoxon *p* |
|---|---|---|---|---|---|---|
| C0 raw vs C2 Dorado 1.1.1 demux | 4,232 | 1,879 | 2.2× | 1.7–3.1× | 8/8 | 0.0078 |
| C1 raw vs C3 Dorado 2.0.0 demux | 4,714 | 2,013 | 2.2× | 1.7–3.3× | 8/8 | 0.0078 |
| C2 Dorado 1.1.1 demux vs C4 Barbell | 1,879 | 33 | 48.3× | 32.5–73.5× | 8/8 | 0.0078 |
| C3 Dorado 2.0.0 demux vs C5 Barbell | 2,013 | 37 | 52.0× | 29.2–70.0× | 8/8 | 0.0078 |
| C2 Dorado 1.1.1 demux vs C3 Dorado 2.0.0 demux | 1,879 | 2,013 | 0.9× | 0.8–1.0× | 0/8 | 0.0078 |

Fold-change is the per-experiment ratio of the first condition over the second, summarized as median (range). p is from the two-sided Wilcoxon signed-rank test (minimum attainable p with n = 8: 0.0078). "Runs A>B" is the number of experiments in which the first condition exceeded the second. The geometric-mean fold-changes, the summary used in the main text, are 2.21× (C0 vs C2), 2.27× (C1 vs C3), 50.0× (C2 vs C4), 50.7× (C3 vs C5), and 0.93× (C2 vs C3). The p-values are not adjusted for multiple comparisons, because the five comparisons address different processing steps rather than repeated tests of one hypothesis; a Bonferroni adjustment for five comparisons would give p = 0.039 for each.

**Table S7.** Reconciliation of strict complete-query HSPs, loci, and distinct barcode-accession pairs in the four query panels.

| Counting stage | B24 | K40 | K46 | K68 |
|---|---|---|---|---|
| (1) Strict complete-query HSPs (both orientations) | 1,390 | 574 | 424 | 6 |
| (2) Barcode/accession/coordinate loci | 695 | 287 | 212 | 3 |
| (3) Distinct barcode-accession pairs | 505 | 224 | 170 | 3 |
| (4) Distinct accessions | 482 | 223 | 170 | 3 |
| (5) Barcodes with matches | 64 | 42 | 41 | 2 |
| (6) Exported TaxID groups | 195 | 114 | 91 | 3 |

All counts use the criterion in S9.3. Each locus is matched once by each of the two query orientations, so row 1 is twice row 2. The panels are nested (K68 within K46 within K40 within B24), so counts are not additive across columns. Figure 2 uses the B24 barcode-accession pairs (row 3). HSP, high-scoring segment pair; TaxID, NCBI Taxonomy identifier.

**Table S8.** Hit-bearing barcodes with a single barcode-only accession, which Figure 2B omits.

| Barcode | Longest construct | Record category | TaxID | Scientific name | Rank |
|---|---|---|---|---|---|
| NB26 | Dorado window (K46) | Plasmid | 1423 | *Bacillus subtilis* | species |
| NB30 | Dorado window (K46) | Organelle | 36190 | *Chaenocephalus aceratus* | species |
| NB37 | Dorado window (K46) | rRNA gene | 5061 | *Aspergillus niger* | species |
| NB50 | Dorado window (K46) | rRNA gene | 5059 | *Aspergillus flavus* | species |
| NB51 | Barcode only (B24) | Plasmid | 1280 | *Staphylococcus aureus* | species |
| NB53 | Dorado window (K46) | Plasmid | 3047468 | *Rhodococcus* sp. AH-ZY2 | species |
| NB55 | Barcode only (B24) | Plasmid | 562 | *Escherichia coli* | species |
| NB58 | Dorado window (K46) | Organelle | 52237 | *Champsocephalus gunnari* | species |
| NB59 | Barcode only (B24) | rRNA gene | 196035 | *Leptopogon amaurocephalus* | species |
| NB62 | Barcode only (B24) | Chromosome/genome | 1423 | *Bacillus subtilis* | species |
| NB67 | Barcode only (B24) | Plasmid | 562 | *Escherichia coli* | species |
| NB70 | Barcode only (B24) | rRNA gene | 4814 | *Chytriomyces hyalinus* | species |
| NB71 | Dorado window (K46) | Plasmid | 1245 | *Leuconostoc mesenteroides* | species |
| NB76 | Barcode only (B24) | Plasmid | 287 | *Pseudomonas aeruginosa* | species |
| NB84 | Barcode only (B24) | Chromosome/genome | 48408 | *Photobacterium leiognathi* subsp. *mandapamensis* | subspecies |
| NB87 | Dorado window (K46) | Plasmid | 562 | *Escherichia coli* | species |
| NB90 | Barcode only (B24) | Chromosome/genome | 3391839 | *Erwinia* sp. LJJL01 | species |
| NB94 | Barcode only (B24) | Plasmid | 562 | *Escherichia coli* | species |
| NB96 | Dorado window (K46) | Phage/virus | 3384911 | Pseudomonas phage BARC01 | species |

Longest construct matched at the barcode locus, record-title category, and exported TaxID with scientific name and rank from NCBI Taxonomy (9 October 2026). Source data: `Fig2B_single_accession_barcodes.csv`. B24, barcode-only query (24 nt); K46, Dorado barcode search window (46 nt); TaxID, NCBI Taxonomy identifier.

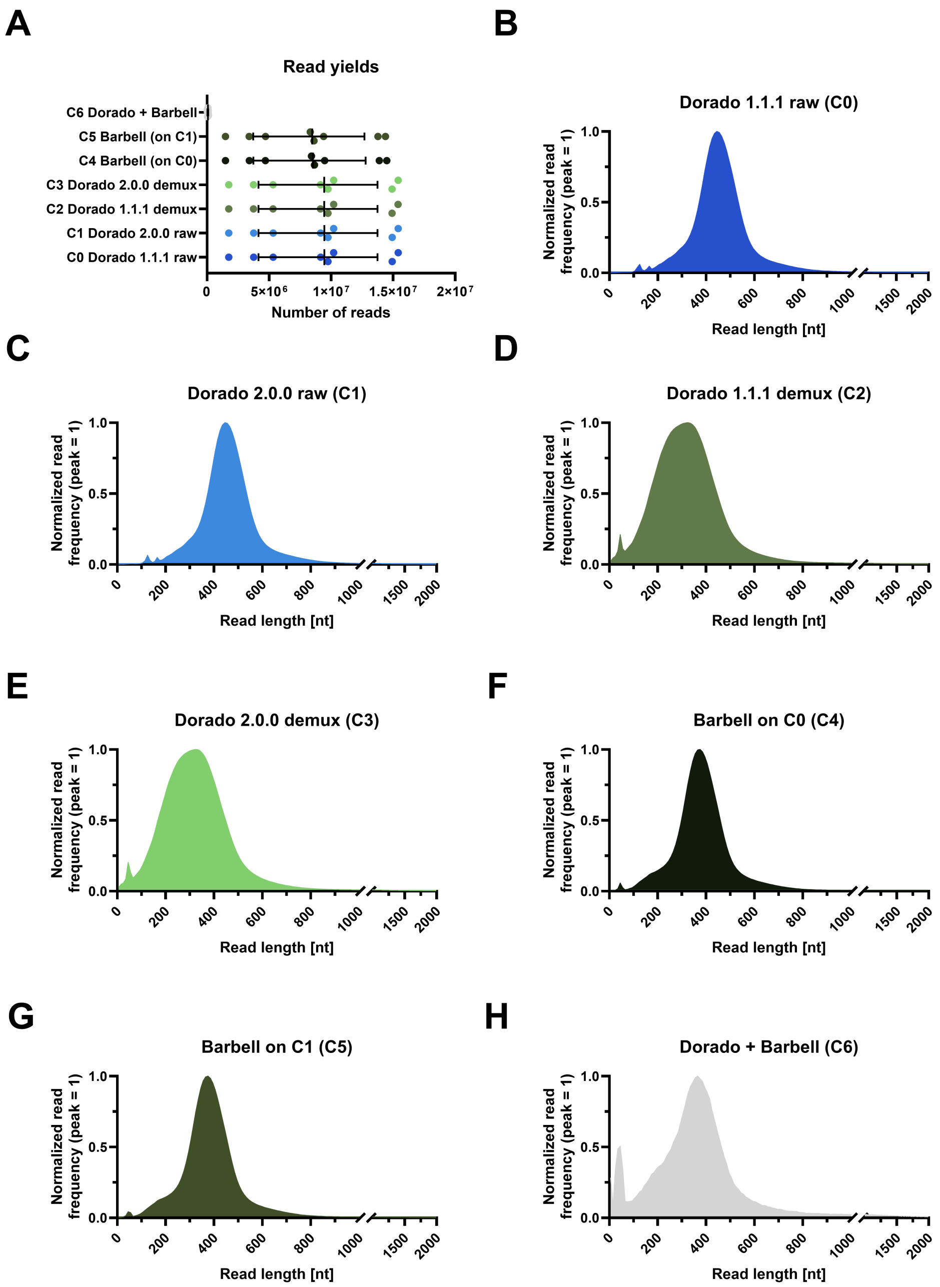


**Figure S1. Read yields and normalized read-length distributions across C0–C6.** C0/C1 are untrimmed Dorado 1.1.1/2.0.0 outputs; C2/C3 are the corresponding Dorado-demultiplexed outputs; C4/C5 apply Barbell directly to C0/C1; C6 applies Barbell to the complete C3 output. (A) Output reads per experiment and condition. Symbols are the eight experiments, lines the median, and whiskers the interquartile range. C6 yields (mean 64,868 ± 32,614 reads per experiment) are close to zero on this linear axis. (B–H) Read-length distributions of C0 (B), C1 (C), C2 (D), C3 (E), C4 (F), C5 (G), and C6 (H). For each condition, output reads were pooled across the eight experiments and counted in 10 nt bins (C0 70,266,179; C1 70,266,869; C2 70,223,803; C3 70,223,900; C4 64,527,600; C5 64,076,165; C6 518,947 reads). Each bin count was divided by the largest bin of the same condition, so every distribution peaks at 1 and the other bins are proportional to it. Filled curves connect consecutive bins. In panels B–H, the x-axis is linear from 0 to 1,000 nt and compressed beyond the axis break; reads of at least 2,000 nt are not shown (1.5–1.9% of C0–C5 and 3.5% of C6 reads). The modal bins were 440–449 nt (C0/C1), 320–329 nt (C2/C3), 370–379 nt (C4/C5), and 360–369 nt (C6). The normalization removes differences in read yield, and the pooled distributions do not show between-experiment variation. nt, nucleotides.
